# Peroxisomal ATPase ATAD1 acts in quality control of the protein import machinery

**DOI:** 10.1101/2022.04.22.489171

**Authors:** Julia Ott, Jessica Sehr, Claudia Lindemann, Katalin Barkovits, Verian Bader, Konstanze Winklhofer, Katrin Marcus, Wolfgang Schliebs, Ralf Erdmann

## Abstract

ATAD1 is an AAA-ATPase which shows a dual localization at mitochondria and peroxisomes. While its peroxisomal function is not known, in mitochondria ATAD1 is part of a quality control mechanism extracting mislocalised tail-anchored and accumulated precursor proteins from the outer membrane. Here, we studied the peroxisomal interactome of ATAD1 and could show that human ATAD1 interacts with PEX5, a cytosolic receptor for peroxisomal matrix proteins which transiently inserts into peroxisomal membranes. Upon cargo-release, mono-ubiquitinated PEX5 is recycled into the cytosol by the AAA-peroxins PEX1 and PEX6. The accumulation of ubiquitinated PEX5 is known to trigger degradation of whole organelles called pexophagy. Here, we used ATAD1-, PEX1- and ATAD1/PEX1-CRISPR-Knockout cell lines to investigate the physiological role of an ATAD1-PEX5 interaction. We could show an influence of ATAD1 on the stability of accumulated PEX5 and hypothesize a role in a peroxisomal quality control mechanism enabling clearance of ubiquitinated receptor from the membrane.

## Introduction

The family of ATPases associated with various cellular activities (AAA family) is characterized by the presence of one or two evolutionary conserved domains containing typical NTP-binding and –hydrolyzing motifs (Walker A and Walker B motifs, N-Linker domain)^1,2^. In line with their broad variety of functions, AAA proteins are located at manifold sites within the cell^3^. In many cases, they attach to membranes to provide mechanic work driven by ATP-hydrolysis, often resulting in disassembly of protein complexes or dislocalization of proteins. Human ATPase family AAA domain-containing protein 1 (ATAD1) like its yeast orthologue Msp1 anchors at mitochondrial and peroxisomal membranes via a helical transmembrane region at the N-terminus thereby exposing its single AAA domain at its C-terminus into the cytosol^4,5^. While the bulk of ATAD1 displays mitochondrial localization in human cells, only a partial association with peroxisomes was indicated^6-9^. In contrast to the mitochondrial function not much is known about the peroxisomal counterpart.

For human ATAD1, and also its yeast homolog Msp1 it has been shown that the protein fulfills a critical role as part of a quality control system of the outer mitochondrial membrane amended for mistargeted tail-anchored (TA) proteins^7-11^. In this respect, Msp1p was shown to act as a dislocase for tail-anchored proteins in a reconstituted system^12^. Recently, another quality-control function of mitochondrial Msp1 had been assigned in the removal of non-imported precursors from translocation pores and mitochondrial surface^11,13^. This inducible surveillance pathway, leading to proteasomal degradation of accumulating precursors was named mitochondrial compromised protein import response (MitoCPR)^13,14^. To induce mitochondrial import stress, Weidberg and Amon overexpressed proteins relying on a bipartite signal sequence and thereby overwhelmed the import capacity leading to accumulation of non-imported precursor proteins at the membrane.

The peroxisomal import system differs from the mitochondrial machinery in several aspects. Most importantly, peroxisomes can translocate folded, even oligomerized proteins^15^. The large and highly flexible import channel is formed by membrane bound import receptors, either PEX5 or one of the PTS2-import co-receptors and their docking protein at the peroxisomal membrane, PEX14^16,17^. PEX5 is a cycling receptor for proteins which possess a peroxisomal targeting sequence type 1 (PTS1) at their C-termini and shuttles between cytosolic and membrane-bound states. Extraction of the receptors from the membrane is facilitated by the AAA+ peroxins PEX1 and PEX6 which use Pex15 in yeast or PEX26 in mammalian cells as an anchor protein at the peroxisomal membrane^18,19^.

To study the function of peroxisome-associated ATAD1 we analyzed the interactome of human ATAD1 at the peroxisomal membrane. Interestingly, among all known peroxisomal membrane proteins we also identified PEX5 as interaction partner. We could demonstrate that in the absence of PEX1 ATAD1 is involved in the removal of accumulating receptor from the membrane.

## Results

### Protein A tagged ATAD1 and inactive ATAD1(E139Q) variant associate with peroxisomes

The aim of this study was to identify potential binding partners of ATAD1 in human cells. For this purpose, Protein A affinity enrichment of ATAD1 was combined with mass spectrometry-based proteomic analysis to enable the identification of ATAD1-interacting proteins. In addition to wild type ATAD1 also the interactome of the hydrolysis inactivating ATAD1(E139Q) variant was investigated, which has been used as a potential substrate trap mutant previously^7^. To this end, ATAD1-TEV-Protein A (ATAD1-TPA) fusion gene was genomically integrated into Flp-In™-293 cells. In a different cell line, we introduced a genomic copy of an ATAD1-TPA fusion gene containing the hydrolysis inactivating mutation E193Q (ATAD1 E193Q-TPA) which has been used as a potential substrate trap mutant previously^7^. As negative control we have used Flp-In293 wild type cells as well as Flp-In293 TEV-Protein A cells (ProtA), where the tag without a fusion gene was genomically integrated.

To ensure the mitochondrial and peroxisomal localization of both ATAD1 variants, we applied immunofluorescence microscopy using anti Protein A antibodies (Supplementary Fig. 1 and Fig. 1). The majority of both Protein A fusion variants are associated with mitochondria. They appeared as filamentous, sometimes ring-shaped structures, which might result from fragmentation of the mitochondrial network (Supplementary Fig. 1). A fraction of ATAD1 appeared in small dots co-localizing with the peroxisomal marker protein PEX14 (Fig. 1A). Consistently with previous observations^6-8^, only a subset of peroxisomes contained ATAD1 in immunodetectable amounts. Overall, neither Protein A fusion nor the E193Q mutation significantly affects the dual peroxisomal and mitochondrial localization of ATAD1 in human cells. This conclusion was further supported by biochemical means. Subcellular fractionation analysis revealed that both Protein A-tagged variants of ATAD1 display same steady-state level and subcellular distribution as endogenous ATAD1 (Fig. 1B). As already indicated in literature^7,8^, both endogenous as well as overexpressed ATAD1 are found in the organellar fraction.

**Fig. 1:**
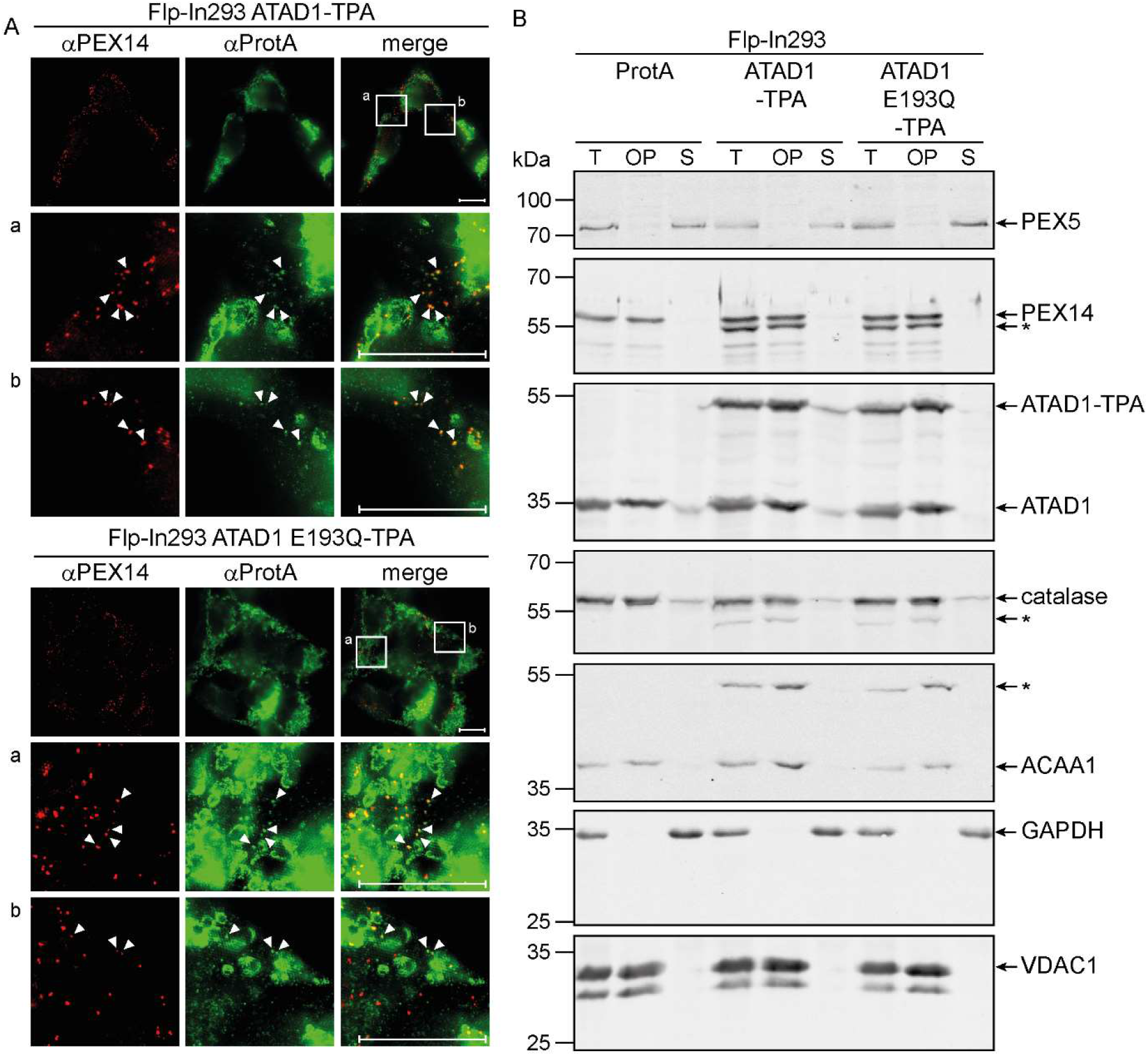
Characterization of the generated cell lines Flp-In293 ATAD1-TPA and ATAD1 E193Q-TPA by immunofluorescence and subcellular fractionation. **A**: Immunofluorescence analysis of Flp-In293 ATAD1-TPA and ATAD1 E193Q-TPA shows that ATAD1 and ATAD1 E193Q colocalize at peroxisomes. Cells were cultivated on coverslips and stained with antibodies against PEX14 and ProtA. The insets (a, b) show magnifications with increased brightness of the ProtA signal for better visibility of punctate ATAD1. Colocalization of ATAD1-TPA and PEX14 are indicated by arrows. Scale bar: 10 µm. **B**: Stable expression of ATAD1-TPA and ATAD1 E193Q-TPA has no influence on the localization of several marker proteins. Flp-In293 ProtA, ATAD1- and ATAD1-E193Q-TPA cells were cultivated on 10 cm plates, analysed by subcellular fractionation and SDS-PAGE followed by immunoblot analysis using the indicated antibodies. Bands marked with an asterisk show cross-reactions of the antibodies with ATAD1-TPA. T: total, OP: organellar pellet, S: supernatant.

In addition, we analyzed the expression level and subcellular localization of several marker proteins in the two generated cell lines (Fig. 1B). The stability of the mitochondrial outer membrane protein VDAC1 is not affected when compared with Protein A-expressing Flp-In293 cells. This suggests that the additional presence of ATAD1 in the same membrane compartment does not induce degradation of mitochondrial membrane proteins via proteasomal or autophagic processes. Furthermore, the majority of peroxisomal matrix enzymes catalase and ACAA1 as well as the peroxisomal receptor docking protein PEX14 sediments in the organellar pellet fraction, independent of additional expression of ATAD1-TPA or ATAD1 E193Q-TPA. This suggests that the import of matrix enzymes into peroxisomes is still functional in both cell lines.

While most peroxisomal marker proteins are behaving similar in the three compared cell lines, we sometimes observed a slight decline in the PEX5 level of the ATAD1-TPA expressing cell line when compared to Protein A and ATAD1 E193Q expressing cells (Fig. 1B). We pursued this observation below.

### ATAD1 interacts with numerous mitochondrial and peroxisomal membrane proteins

Prior to the proteomic analysis of ATAD1 and ATAD1 E193Q complexes, the expression and isolation of the ATAD1-TPA variants as well as the enrichment of the complexes after affinity purification and subsequent elution by TEV protease cleavage were investigated (Fig. 2). Therefore, differential centrifugation and solubilisation were performed to generate distinct subtractions of the corresponding cell lysates (Fig. 2A). Western blot analysis using ATAD1-detecting antibodies confirmed the expression and enrichment of both ATAD1 variants and further revealed that the steady-state level of ATAD1 (E193Q)-TPA in each cell line was similar compared with the expression of endogenous one (Fig. 2B/C). After affinity purification followed by TEV protease cleavage, recombinant as well as endogenous protein could be detected for the ATAD1 and the ATAD1 E193Q complex (Fig. 2B/C and Supplementary Fig. 2B). Coomassie-staining of the TEV-eluate fractions revealed the enrichment of several proteins by affinity purification of ATAD1 (E193Q)-TPA complexes (Supplementary Fig. 2A). Given that in the negative control (Flp-In293 cells lacking TPA-labelled ATAD1) endogenous ATAD1 was not immunologically detectable after affinity purification in the TEV protease-cleaved eluate, affinity co-purification of the ectopically expressed variant with approximately equimolar amounts of endogenous ATAD1 suggests formation of oligomeric, most likely hexameric complexes (Figure 2B/C and Supplementary Fig. 2B).

**Fig. 2:**
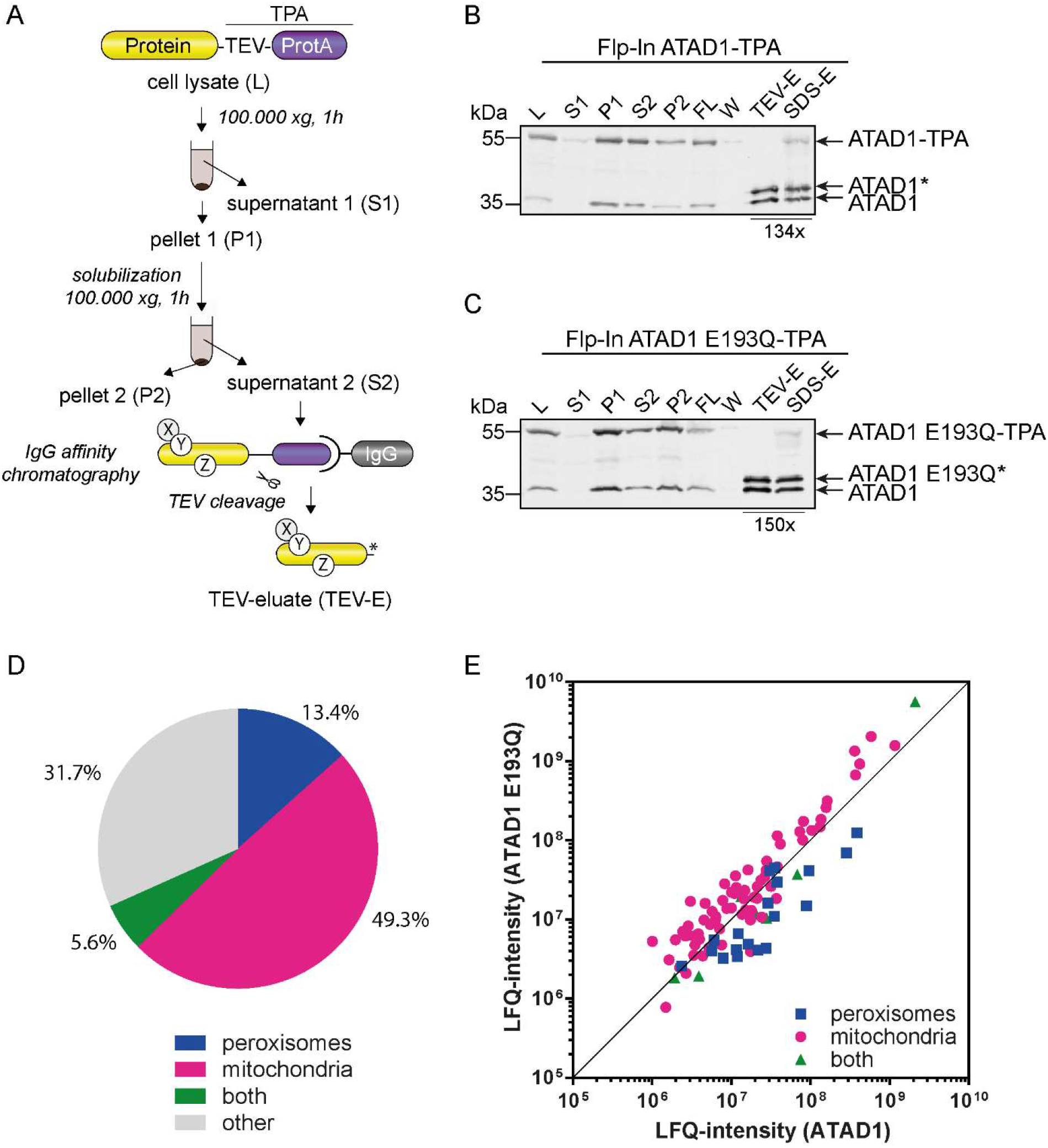
Mass spectrometric analysis of ATAD1 (E193Q)-TPA complexes revealed the interaction to several mitochondrial and peroxisomal proteins. **A:** Cell lysates (L) of TPA-fusion constructs expressing cells were prepared by disrupting of 5 g harvested cells with glass beads. Soluble (S1) and membrane-bound complexes (P1) were separated by ultracentrifugation (100.000xg, 1 h, 4°C). After solubilization of membrane complexes with 1% digitonin (w/v) non-soluble components were removed by centrifugation at 100.000xg for 1 h (P2), while the solubilized membrane complexes (S2) were incubated with IgG-sepharose overnight. After removal of non-bound complexes by spinning and washing of the sepharose complexes were eluted by TEV-cleavage (2 h, 16°C). **B/C:** Immunoblot analysis of SDS-samples using a primary ATAD1-antibody after each preparation step. Besides TEV-cleaved ATAD1 which is denoted with an asterisk (*) also endogenous protein could be detected in the eluates. **D:** Most of the proteins which fulfilled the criteria for being identified as a specific binding partner in ATAD1 (E193Q)-TPA complexes were mitochondrial proteins, followed by peroxisomal proteins. **E:** Mitochondrial proteins were enriched in ATAD1 E193Q -TPA complexes, while peroxisomal proteins were enriched in ATAD1-TPA complexes. To visualize the distribution of mitochondrial and peroxisomal proteins within ATAD1 (E193Q)-TPA complexes the LFQ-intensity in which they appeared in each complex was plotted. Diagonal line shows equal appearance within wild type and mutated complex.

After successful verification of affinity-based purification of the two ATAD1 complexes, five independent purification approaches for each ATAD1 variant and two replicates of a negative control were subjected to proteomic analysis to identify ATAD1-binding partners. In total 3043 proteins were identified in the whole data set. To determine potential binding partners and exclude nonspecific proteins, strict filter criteria were applied, including the exclusive selection of proteins that were more than 5-fold enriched in the respective ATAD1 purification approach compared with the negative control and were furthermore identified in at least 4 replicates out of 5. With this, 142 proteins were selected as potential ATAD1-binding partners (Supplementary Tab. 1). Among these we found 19 membrane proteins exclusively associated with peroxisomes and 8 membrane proteins located to both peroxisomes and mitochondria (Tab. 1) which can be divided into the three functional groups peroxins, transporter and enzymes. Besides the TA-proteins PEX26, FAR1, FAR2, MFF and MAVS we found every single type of membrane proteins (I-V), which confirms the already made observation that the substrate specificity of ATAD1 is not limited on TA-proteins^13^.

**Table 1:** Peroxisomal proteins found in ATAD1 (E193Q)-TPA complexes by mass spectrometric analysis. Peroxisomal proteins were highly enriched in the affinity purified ATAD1 complexes compared to negative control complexes from wild type cells. To show enrichment in wild type (ATAD1) or mutated (ATAD1 E193Q) complex the enrichment factor was calculated by dividing the LFQ-intensity through the negative control (LFQ ATAD1(E193Q)/LFQ control). Proteins are sorted descending by their enrichment in the ATAD1 complex. Blue: peroxisomal proteins, green: both peroxisomal and mitochondrial proteins. Proteins marked with an asterisk were not observed in the negative control. Abbreviations: TM: transmembrane; ss: single-span; ms: multi-span; TA: tail-anchored; C: cytosol; PA: peripherally attached; M: matrix.

| protein | protein name | enrichment factor |  | topology |
| --- | --- | --- | --- | --- |
|  |  | ATAD1 | ATAD1 E193Q |  |
| PEX11B | Peroxisomal membrane protein 11B | 73.4 | 31.7 | ss TM |
| PXMP2 | Peroxisomal membrane protein 2 | 56.5 | 9.4 | ms TM |
| PEX3 | Peroxisomal biogenesis factor 3 | 39.9 | 12.5 | ms TM |
| SLC25A17 | Peroxisomal membrane protein PMP34 | 37.5 | 7.1 | ms TM |
| PEX11G | Peroxisomal membrane protein 11C | 23.5 | 16.5 | ms TM |
| PEX19 | Peroxisomal biogenesis factor 19 | 19.8 | 15.5 | C/PA |
| PEX16 | Peroxisomal membrane protein 16 | 19.0 | 10.3 | ms TM |
| PEX10 | Peroxisome biogenesis factor 10 | 18.6 | 6.7 | ms TM |
| ABCD3 | ATP-binding cassette sub-family D member 3 | 16.7 | 5.4 | ms TM |
| PEX2 | Peroxisome biogenesis factor 2 | 14.0 | 4.0 | ms TM |
| PEX5 | Peroxisomal targeting signal 1 receptor | 12.1 | 3.6 | C/PA |
| FAR2 | Fatty acyl-CoA reductase 2 | 9.8 | 1.5 | TA |
| FAR1 | Fatty acyl-CoA reductase 1 | 9.7 | 2.3 | TA |
| PEX6 | Peroxisome assembly factor 2 | 6.3 | 7.9 | PA (C) |
| PEX1 | Peroxisome biogenesis factor 1 | 5.5 | 7.4 | PA (C) |
| GNPAT | Dihydroxyacetone phosphate acyltransferase | 5.5 | 3.0 | PA (M) |
| PEX12 | Peroxisome assembly protein 12 | * | * | ms TM |
| PXMP4 | Peroxisomal membrane protein 4 | * | * | ms TM |
| PEX26 | Peroxisome assembly protein 26 | * | * | TA |
| ATAD1 | ATPase family AAA domain-containing protein 1 | 211.0 | 562.2 | ss TM |
| MFF | Mitochondrial fission factor | 49.1 | 26.6 | TA |
| MARC2 | MOSC domain-containing protein 2, mitochondrial | 22.4 | 36.4 | ? |
| ABCD1 | ATP-binding cassette sub-family D member 1 | 6.5 | 2.4 | ms TM |
| ACSL1 | Long-chain-fatty-acid--CoA ligase 1 | 5.2 | 3.0 | ss TM |
| MAVS | Mitochondrial antiviral-signaling protein | 4.1 | 6.0 | TA |
| TMEM135 | Transmembrane protein 135 | * | * | ms TM |
| MUL1 | Mitochondrial ubiquitin ligase activator of NFKB 1 | * | * | ms TM |

There are two known TA proteins which act as targets of ATAD1 when mislocalised to the outer membrane of mitochondria: PEX26 and endosomal GOSR1^7^. We found both in the complex but only PEX26 fulfilled the criteria for being selected as a specific binding partner. The Golgi SNAP receptor complex member 1 (GOSR1) was only present in the trapping mutant ATAD1 E193Q and not in the ATAD1 complex which is consistent with the literature^7^, but it was not 5-fold enriched compared to the negative control. In contrast, the TA peroxisomal PEX26 (yeast Pex15p homolog) which has been characterized as the archetype of ATAD1 substrates when mislocalized to the outer membrane of mitochondria^7,8,10^ was found in both ATAD1 and ATAD1 E193Q complex in comparable amounts. This means it was not enriched in association with the trapping ATAD1 mutant compared to its abundance in ATAD1 eluates (LQF intensity [ATAD1 E193Q/ATAD1] ratio of 1.09, see Supplementary Tab. 1), which was unexpected.

While we did not find all known interaction partners of ATAD1 in ATAD1 or ATAD1 E193Q complexes, more than 90 % of the identified proteins which fulfilled all criteria were membrane proteins or at least partially attached to membranes (Supplementary Tab. 1). Most of them are localized to mitochondria and peroxisomes. Interestingly, almost all identified peroxisomal membrane proteins as shown in Fig. 2E exhibit stronger binding to the non-mutated ATAD1 than to ATAD1 E193Q (Tab. 1, Supplementary Tab. 1), while for mitochondrial proteins it behaves exactly the other way (Supplementary Tab. 2). In contrast to mitochondria interaction, the capability of ATP-hydrolysis seems to be advantageous for ATAD1 association with peroxisomal membrane proteins. This suggests that ATP-derived energy is used for an unknown process preceding protein-protein interaction at the peroxisomal membranes.

Among the 142 proteins from Supplementary Tab. 1 we found nearly all members of the peroxisomal import machinery together with other PMPs summarized in Tab. 1. Two proteins particularly catch the eye, as they are mostly cytosolic proteins and only in a small amount localized to the peroxisomal membrane: PEX5 and PEX19. While PEX19 at least has a farnesylation site which could assist in membrane targeting^20,21^, PEX5 is lacking any known membrane domains. This has drawn our attention on PEX5, as its properties differ from the known ATAD1 targets, which exclusively are membrane proteins with no cytosolic localization^7,13^.

### ATAD1 facilitates degradation of PEX5 in PEX1-deficient cells

Maintaining protein homeostasis is one of the crucial mechanisms of the cell to sustain cellular processes. For mitochondria there are several quality control mechanisms known which use the dislocation ability of ATAD1 to achieve this^11^, one of which is clearing stalled precursor proteins from the mitochondrial translocation machinery^13^. In line with this we could identify the peroxisomal import receptor PEX5 as an interaction partner of ATAD1 by affinity purification. This raised the question whether ATAD1 acts in a quality control mechanism at the peroxisomal import pore similar to mitochondria (peroxiCPR). One possibility to address this was to disturb the protein homeostasis of peroxisomes and trigger currently unknown quality control mechanisms. Therefore, we created a situation where the export of PEX5 from the peroxisomal membrane is impeded by deletion of PEX1 (Fig. 3A/B). This was done by the generation of several T-REx293 KO-cell lines using the CRISPR/Cas9 technique (not published yet, will be described elsewhere). The preparation and western blot analysis of whole cell lysates of these KO cell lines indeed showed ATAD1-dependent differences in the steady-state level of PEX5 which will be describe below (Fig. 3C).

**Fig. 3:**
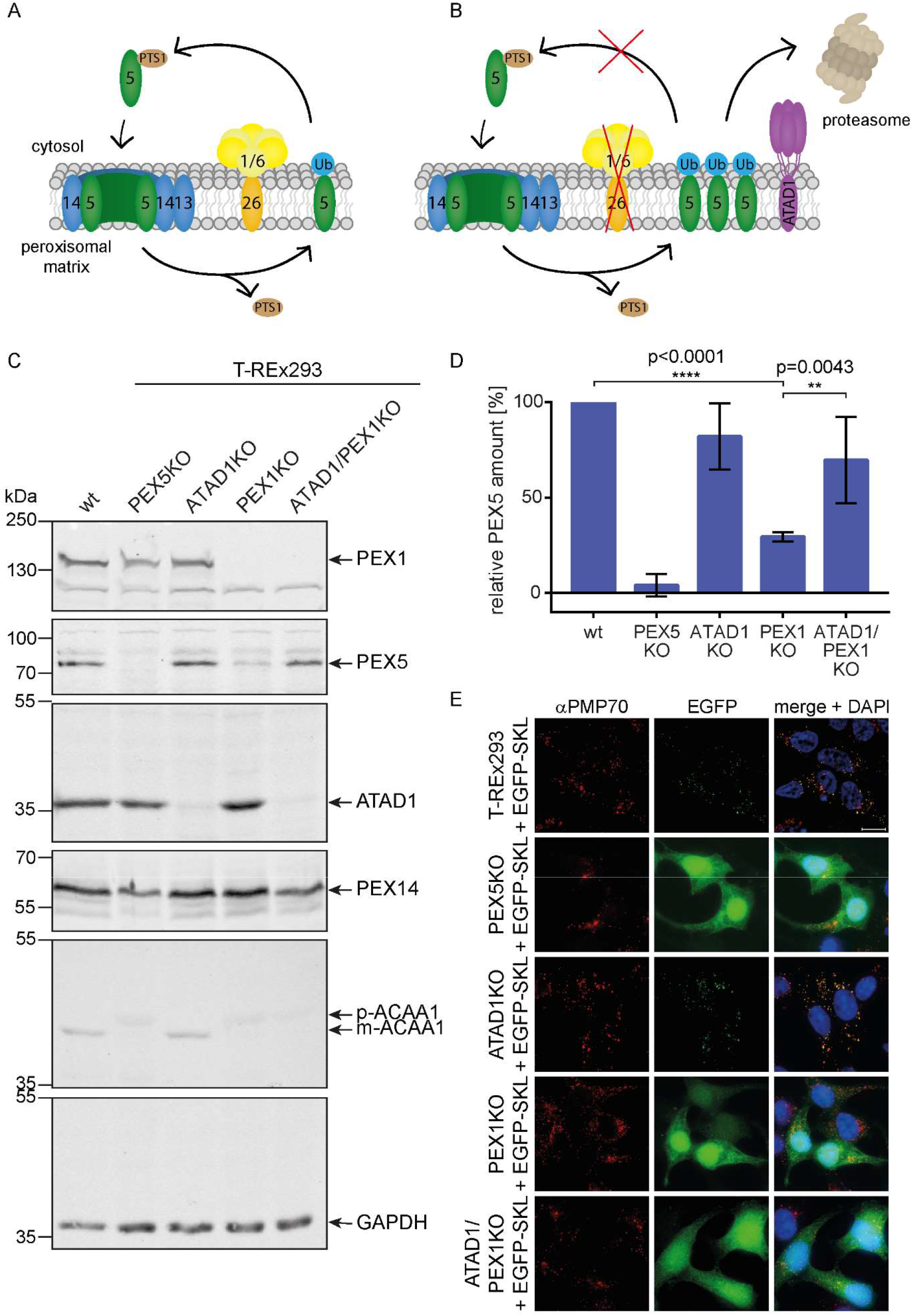
ATAD1 shows an influence on PEX5-stability in case of PEX1-deletion. **A:** Functional peroxisomal matrix protein import cycle. PEX5 is mono-ubiquitinated after releasing the cargo into the peroxisomal matrix and extracted by the export complex consisting of PEX1/PEX6/PEX26. **B:** Proposed model for the peroxisomal quality control mechanism peroxiCPR. An impaired recycling of the peroxisomal receptor PEX5 due to PEX1-deletion leads to its accumulation at the peroxisomal membrane. To limit the damage by avoiding pexophagy which would be triggered by the accumulation of ubiquitinated receptor, PEX5 is recognized by the dislocase ATAD1 and degraded by the proteasome. **C:** Whole cell lysates of T-REx293 wild type, PEX5KO, ATAD1KO, PEX1KO and ATAD1/PEX1KO cells were prepared and analyzed by SDS-PAGE and immunoblot analysis with the indicated antibodies. A deletion of PEX1 causes degradation of PEX5, while an additional ATAD1-deletion stabilizes PEX5 again. **D:** Quantification of the PEX5 amount in KO cell lines. PEX5 amount in wt cells was set as 100%. Data represent mean ± SD. PEX5-stability in ATAD1/PEX1KO is significantly increased compared to PEX1KO cells as analyzed by one-way ANOVA, n=4. **E:** The reduced stability of PEX5 in the PEX1KO cell line is not caused by pexophagy. Cells were cultivated on coverslips and transfected with EGFP-SKL. 48 h after transfection cells were fixed with formaldehyde and PMP70 was visualized with a PMP70 antibody. Wild type and ATAD1KO cells show a punctated pattern of EGFP-SKL which colocalizes with PMP70, while PEX5KO, PEX1KO and ATAD1/PEX1KO cells exhibit cytosolically mislocalized EGFP-SKL. Scale bar: 10 µm.

Despite a functional peroxisomal export complex a single deletion of ATAD1 has no influence on the stability of the receptor PEX5. An impeded export of PEX5 due to a non-functional export machinery however would lead to accumulation of the receptor at the peroxisomal membrane^22^. In PBD-patient fibroblast cell lines it has already been shown that the absence of functional PEX1 causes a reduced PEX5 stability due to proteasomal degradation^23^. We could show that a deletion of PEX1 leads to a reduced steady-state level of PEX5 to about 30% of wild type level (Fig. 3D). Interestingly, a double deletion of ATAD1 and PEX1 shows a stabilization of PEX5 compared to a single PEX1KO (Fig. 3C). The PEX5-amount in ATAD1/PEX1KO cells reaches about 70% of wild type level (Fig. 3D). This result shows that ATAD1 is responsible for the reduced PEX5-stability in PEX1 deficient cells.

It has been shown that a loss of the peroxisomal export complex leads to increased pexophagy, which would be a possible explanation for the reduced PEX5 stability in PEX1KO cells^22,24^. However, in our case we don’t see a reduced level of peroxisomes in PEX1KO cells compared to wild type cells (Fig. 3E), so we propose a primary role of the proteasome in reducing the stability of PEX5.

## Discussion

Here, we discovered a function for ATAD1 in the quality control of peroxisome protein import that resembles in several aspects the mitoCPR strategy. MitoCPR was recently detected as a surveillance mechanism which antagonizes severe defects in mitochondrial protein import by clearing stalled precursor proteins from the translocation machinery^13^. The central constituent of the peroxisomal import channel is the cycling receptor PEX5 which is extracted from the membrane by the AAA ATPases complex. In human T-REx293 cells, we knocked out PEX1, which is one of the ATPases, to disturb the peroxisomal protein homeostasis using CRISPR/Cas9. This led to a reduced PEX5-stability to about 30% of wild type level. In the literature it was reported that in PEX1 deficient human fibroblasts PEX5 accumulates at the peroxisomal membranes^22,23^. In addition, the stability of cytosolic PEX5 was severely reduced (about 20-fold) in cells defective in recycling of the receptor, which is supported by our data. It is suggestive, that an unknown machinery removes the accumulated receptor from the peroxisomal membrane leading to degradation, most likely through the proteasome. We could not exclude the possibility that PEX5 abundance is reduced in these cells due to pexophagic degradation of organelles accumulating ubiquitinated receptor^22^. However, we do not see a reduction of peroxisome number or abundance of peroxisomal marker proteins in PEX1 KO cells when comparing with wild type (Fig. 3) supporting the view that stacked PEX5 is specifically extracted and degraded from the membrane. One of the proteins which is responsible for the instability of PEX5 in recycling-deficient cells is ATAD1 as clearly indicated by the double KO which shows a wild type like abundance of the receptor (Fig. 3).

However, KO of ATAD1 could not recover PEX5-mediated protein import into peroxisomes in PEX1 deficient cells, despite a significant increase of the steady-state level of the receptor PEX5. Though the question arises what might be the function of the ATAD1-dependent rescue of PEX5. We think that the major function of peroxiCPR is to remove ubiquitinated PEX5 to avoid degradation of metabolically active peroxisomes via pexophagy^22^. To this end, it is important to note that in mitochondria several quality control systems exist in order to suppress functional damage and mitophagy as a response to protein import stress. Besides ATAD1-dependent mitoCPR, deficits in protein import trigger the unfolded protein response (mtUPR) which leads to transcriptional induction of various protective activities located inside mitochondria, i.e. chaperones and proteases. Recently identified unfolded protein response activated by protein mistargeting (UPRam) links accumulation of precursor proteins in the cytosol with the activation of proteasomal activities^25^. Whether such import stress sensitive pathways as found with mitochondrial import defects also exist for defects in peroxisome biogenesis is a matter of debate.

One of the central questions of our study concerns the peroxisomal substrate specificity of ATAD1. Originally it was believed that mistargeted tail-anchored proteins are the only cognate substrates. But the involvement of ATAD1 in mitoCPR as well as peroxiCPR demonstrates that neither mislocalization nor the topology of membrane proteins alone define specificity. This is further supported by our proteomic analysis which shows that almost all peroxisomal membrane proteins (at least after detergent solubilization) associate with ATAD1 (Tab. 1). No specificity for tail-anchored proteins was indicated. It seems that ranking of detergent solved PMPs depends on abundance so that every PMP can bind. Same is valid for mitochondrial substrates (see Supplementary Tab. 2). Weir et al. could show that overexpressed Pex15 is removed from the peroxisomal membrane by Msp1 in yeast cells. The authors concluded that Pex15 turnover by Msp1 is triggered by the solitary existence of the membrane protein. Pex3 was identified as kind of heteromeric shield of Pex15 preventing its extraction^12^. Solitary existence caused by lack of partner proteins is a common feature of mistargeted but also of accumulated membrane proteins. In non-disturbed peroxisomal membranes PEX5 binds in a 1 to 1 stoichiometry with PEX14. Since the amount of PEX14 is limited and PEX5 enters the peroxisomal docking site steadily without being extracted again, solitary PEX5 will accumulate in the case of import stress. This might explain that ATAD1 is not involved in the regular receptor cycle but becomes important later upon accumulation. In terms of a suitable therapy against peroxisomal biogenesis disorders the overexpression of ATAD1 preventing degradation of peroxisomes could be beneficial. A treatment by increasing the ATAD1 level has been recently considered as kind of therapy for Zellweger patients in which peroxins are mistargeted to mitochondria and perturb their function^26^.

## Experimental Procedures

### Plasmids

For the stable integration of Protein A in Flp-In293 cells the vector pcDNA5/FRT-ATG-TEV-Protein A^27^ was used. The plasmid for the stable integration of ATAD1-TPA was constructed by PCR-amplification of ATAD1 from pCMV-SPORT6-*Hs*ATAD1 (RZPD/ImaGenes, Berlin) with the primers 5’-gatcgatatcatggtacatgctgaagccttt-3’ (for) and 5’-gatcaagcttatctaaacaaacatgtgttaaaac (rev) and inserted into the *EcoRV*/*HindIII* digested vector pcDNA5/FRT-ATG-TPA^27^. The Walker B mutation E193Q was then inserted via Site Directed Mutagenesis with the primer pair 5’-gctacaaccatccatcatctttatagatcaaatagactcctttctacg-3’ (for) and 5’-cgtagaaaggagtctatttgatctataaagatgatggatggttgtagc-3’ (rev). For generation of the CRISPR KO cell lines the following oligonucleotide pairs were annealed and inserted into the pX458 vector (Addgene 48138, Dr. Feng Zhang) to act as guide RNAs: ATAD1 (for 5’-ccgactcaaaggacgagaaa-3’, rev 5’-tttctcgtcctttgagtcgg-3’), PEX1 (for 5’-agcgatgcgct ggcgggtgc-3’, rev 5’-gcacccgccagcgcatcgct-3’), PEX5 (for 5’-cgggttggcacccccgcatt-3’, rev 5’-aatgcgggggtgccaacccg-3’). The generation of the plasmid EGFP-SKL was described in Neuhaus et al.^28^.

### Cell culture

Human embryonic kidney 293 cells (HEK293), Flp-In™-293 and T-REx™-293 cells (Invitrogen, USA) were grown in Dulbecco’s Modified Eagle’s Medium high glucose (DMEM) supplemented with 10 % fetal calf serum, 4 mM L-glutamine, 100,000 U/l penicillin and 100 mg/l streptomycin at 37°C and 8.5% CO_2_. The cell lines Flp-In293 ATAD1-TEV-Protein A (TPA) and ATAD1 E193Q-TPA were generated by genomic integration of ATAD1-TPA and ATAD1 E193Q-TPA fusion genes into Flp-In293 cells^27^. The cell lines T-REx-293 PEX5KO, ATAD1KO, PEX1KO and ATAD1/PEX1KO were generated using the CRISPR/Cas9 system in T-REx-293 cells (not published yet, will be described elsewhere).

For all cell transfections X-tremeGENE HP DNA Transfection Reagent (Roche, Germany) was used according to the manufacturer’s instructions.

### Preparation of whole cell lysates, subcellular fractionation and immunoblot analysis

For the preparation of whole cell lysates cells were seeded to >90% confluence on 6 cm dishes. After washing with Dulbecco’s Phosphate buffered saline (D’PBS, Gibco, UK) cells were scraped from the dishes, absorbed in D’PBS and harvested 7 min at 200xg and 4°C. The cell sediment was resuspended in lysis buffer (CAT ELISA, Roche, Germany) supplemented with cOmplete™ EDTA-free Protease Inhibitor Cocktail (Roche) and incubated 30 min at 4°C. After sedimentation of cell debris (13,000 rpm, 5 min) SDS-sample buffer was added to the supernatant and samples were analyzed by SDS-PAGE and immunoblot analysis. Band intensities on immunoblots were quantified using densitometry (ImageJ, NHI).

For subcellular fractionation cells were seeded to >90% confluence on 10 cm dishes, washed with HBSS (Hanks Balanced Salt Solution), trypsinized and resuspended in HBSS+MgCl_2_. Cell suspension was divided into two parts (for total fraction and organellar pellet/ soluble fraction) and harvested 7 min at 200xg and 4°C. For the total fraction one cell pellet was resuspended in RIPA buffer (Sigma-Aldrich, Germany) supplemented with protease inhibitors and incubated 30 min at 4°C. After centrifugation at 8,500 rpm for 5 min at 4°C SDS-sample buffer was added to the supernatant. For separation of the organellar and soluble fraction the other cell pellet was resuspended in 0.05 % digitonin in D’PBS supplemented with protease inhibitors and incubated 30 min at 4°C. The organellar pellet was sedimented by centrifugation at 8,500 rpm for 5 min at 4°C. For the soluble fraction SDS-sample buffer was added to the supernatant, while the organellar pellet was resuspended in RIPA buffer supplemented with protease inhibitors and incubated 30 min at 4°C. After centrifugation at 8,500 rpm for 5 min at 4°C SDS-sample buffer was added to the supernatant and samples were analyzed by SDS-PAGE and immunoblot analysis.

Protein level was determined using the following primary antibodies: rabbit aPEX5 (1:5000,^29^), mouse aPEX1 (1:250, BD Biosciences, USA), mouse aATAD1 (1:1000, OriGene, USA), rabbit aPEX14 (1:5000,^30^), mouse aGAPDH (1:7500, Proteintech, Germany), rabbit aACAA1 (1:1000, Sigma-Aldrich, Germany), mouse aCatalase (1:500, Abcam, UK) and mouse aVDAC1 (1:5000, Abcam, UK). As fluorescent secondary antibodies goat a-rabbit and goat a-mouse IRDye 800CW (1:15,000, LI-COR Biosciences, Germany) were used and detected by the Odyssey Infrared Imaging System (LI-COR Biosciences).

### IgG affinity purification

For complex isolation Flp-In293 ATAD1-TPA and ATAD1 E193Q-TPA cells were cultured on triple flasks (500 cm^2^, Nunc, Germany) under normal growth conditions followed by harvesting. 5 g cells (wet weight) were disrupted with glass beads in HEPES lysis buffer supplemented with protease inhibitors. After removal of cell debris (1,500xg, 5 min, 4°C, Eppendorf 5810R) soluble complexes and membrane-bound complexes were separated by ultracentrifugation (100,000xg, 1 h, 4°C, SW41 Ti). Membrane complexes were then solubilized with 1% digitonin (w/v) for 1 h and non-soluble components removed by centrifugation at 100,000xg for 1 h (SW41 Ti). Solubilized membrane-bound complexes were incubated overnight with IgG-sepharose and washed 10 times with washing buffer (0,1% Digitonin (w/v)) by spinning the samples for 1 min at 150xg and 4°C (Centrifuge 5417R). Bound TPA-complexes were eluted by incubation with TEV-protease for 2 h at 16°C and spinning (1 min, 150xg, 4°C, Centrifuge 5417R).

### Mass spectrometry for the identification of ATAD1 interaction partners

For in-gel digestion 4 µg of each affinity purification including ATAD1-/ATAD1 E193Q-complexes as well as bands were hashed and destained by three times alternating 10 min treatments with buffer A (10 mM ammoniumhydrogencarbonate, pH 8.3) and buffer B (buffer A: 100% acetonitrile from Merck KGaA, Darmstadt, Germanyin a ratio of 50:50 (v/v)). After the second incubation with 50 mM ammonium bicarbonate, samples were treated with 50 µl 10 mM DTT (AppliChem GmbH, Darmstadt, Germany) for 1 h at 56 °C and with 50 µl 50 mM IAA (Merck KGaA) for 45 min at room temperature before the destaining protocol was continued. Finally, gel pieces were dried in a vacuum concentrator (RVC2-25CD plus, Martin Christ Gefriertrocknungsanlagen, Osterode am Harz, Germany). Digestion was initiated by adding 8 µl of trypsin solution (0.015 µg/µl, Serva, Heidelberg, Germany) and was performed overnight. The digestion was stopped, and peptides eluted by incubating the gel pieces two times for 15 min with 30 µl of a 1:1 solution containing 100% acetonitrile and 0.1% (v/v) TFA (Merck KGaA, Darmstadt, Germany) in an ice-cooled ultrasonic bath. Samples were dried in a vacuum concentrator and resuspended in 40 µl 0.1% (v/v) TFA. Afterwards, the peptide concentration was determined by amino acid analysis (AAA) as described previously^31^.

Afterwards, 200 ng tryptic peptides were measured by nanoLC-ESI-MS/MS. An UltiMate 3000 RSLC nano LC system (Thermo Scientific, Bremen, Germany) was utilized for nano HPLC analysis using the following solvent system: (A) 0.1% FA; (B) 84% ACN, 0.1% FA. Samples were initially loaded on a trap column (Thermo, 100 μm × 2 cm, particle size 5 μ m, pore size 100 Å, C18) with a flow rate of 30 μl/min with 0.1% TFA. After sample concentration and washing, the trap column was serially connected with an analytical C18 column (Thermo, 75 μ m × 50 cm, particle size 2 μm, pore size 100 Å), and the peptides were separated with a flow rate of 400 nl/min using a solvent gradient of 4% to 40% B for 95 min at 60 °C. After each sample measurement, 1 h of column washing was performed for equilibration. The HPLC system was on-line connected to the nano-electrospray ionization source of a Q-Exactive classic (Thermo Scientific, Bremen, Germany). Full MS spectra were scanned between 350–1400 m/z with a resolution of 70,000 at 200 m/z (AGC target 3e6, 80 ms maximum injection time). Capillary temperature was set to 275 °C and spray voltage to 1500 V (positive mode). Lock mass polydimethylcyclosiloxane (m/z 445.120) was used for internal recalibration. The m/z values initiating MS/MS were set on a dynamic exclusion list for 30 s and the 10 most intensive ions (charge 2+ to 3+) were selected for fragmentation. MS/MS fragments were generated by higher-energy-collision-induced dissociation and the fragmentation was performed with 28% normalized collision energy. The fragment analysis was performed in an orbitrap analyzer with resolution 35,000 at 200 m/z (AGC 2e5, maximum injection time 120 ms).

Peptides and proteins were identified and quantified using MaxQuant (v.1.5.0.0, https://maxquant.org/) using Andromeda as a search engine. Spectra were matched against UniProt/Swiss-Prot using human taxonomy (released 2014_11) with the addition of the ATAD1-E193Q mutant information. Methionine oxidation was set as variable modifications; cysteine carbami-domethylation as a fixed one. The minimum number of peptides and razor peptides for protein identification was 1; the minimum number of unique peptides was 0. Protein identification was performed at a protein false discovery rate of 0.01. The “match between runs” option was on. Label-free quantification was carried out based on LFQ^32^. MaxQuant data analysis and visualization were performed using Microsoft Excel (Microsoft Corporation, USA) and GraphPad Prism (Graphpad Software, Inc, USA).

### Fluorescence microscopy

For immunofluorescence microscopy cells were seeded on coverslips to appropriate density and fixed with 3% formaldehyde in D’PBS for 20 min. For membrane permeabilization cells were incubated in 1% Triton X-100 for 5 min. After blocking with 5% FCS and 2% BSA in D’PBS cells were incubated for 30 min in the primary antibodies chicken aPEX14 (1:500, Ruhr-University Bochum), mouse aTOM20 (1:200, Santa Cruz biotechnology), rabbit aPMP70 (1:500, Invitrogen, USA) and rabbit aProtein A (1:1000, Sigma-Aldrich, Germany). The incubation with the following secondary antibodies (1:300) was done for 10 min protected from light: goat a-rabbit IgG (H+L) Alexa Fluor 488 and 594 (Invitrogen), goat a-mouse IgG (H+L) Alexa Fluor 488 (Invitrogen) and goat a-chicken IgG (H+L) Alexa Fluor 594 (Invitrogen). Coverslips were mounted on glass slides using Mowiol 4-88 (Calbiochem, USA) supplemented with DAPI. Imaging was performed using the Zeiss ELYRA PS.1 Super-Resolution Structured Illumination Microscope (SR-SIM, Zeiss, Germany).

### Statistical analysis

Statistical analysis was performed by one-way ANOVA with Tukey’s multiple comparisons using GraphPad Prism (Graphpad Software, Inc, USA). Data were considered significant at a value of *p < 0.05.

## Abbreviations

AAA: ATPases associated with various cellular activities
ATAD1: ATPase family AAA domain-containing protein 1
CRISPR: clustered regularly interspaced short palindromic repeats
mitoCPR: mitochondrial compromised protein import response
mtUPR: mitochondrial unfolded protein response
PEX: peroxin
PMP: peroxisomal membrane protein
ProtA: Protein A
PTS: peroxisomal targeting sequence
TA: tail-anchored
TPA: TEV-Protein A
UPRam: unfolded protein response activated by protein mistargeting
VDAC1: voltage-dependent anion-selective channel protein 1

