## Supplement for "Peroxisomal ATPase ATAD1 acts in quality control of the protein import machinery"

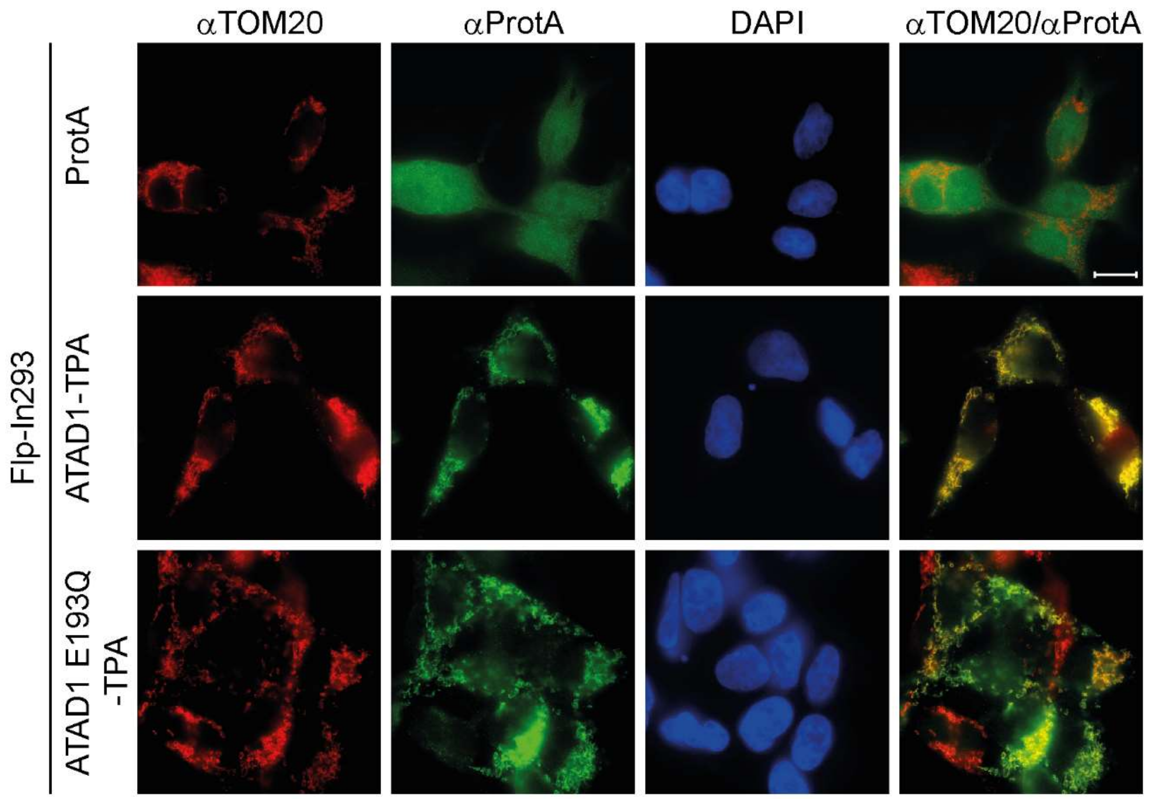


**Supplementary Fig. 1: Immunofluorescence microscopic analysis of Flp-In293 ProtA, ATAD1-TPA and ATAD1 E193Q-TPA cells.** Cells were cultivated on coverslips and stained with antibodies against TOM20 and ProtA. Nuclei were visualized with DAPI. Most of ATAD1-/ATAD1 E193Q-TPA show a colocalization with TOM20. Scale bar: 10 µm.

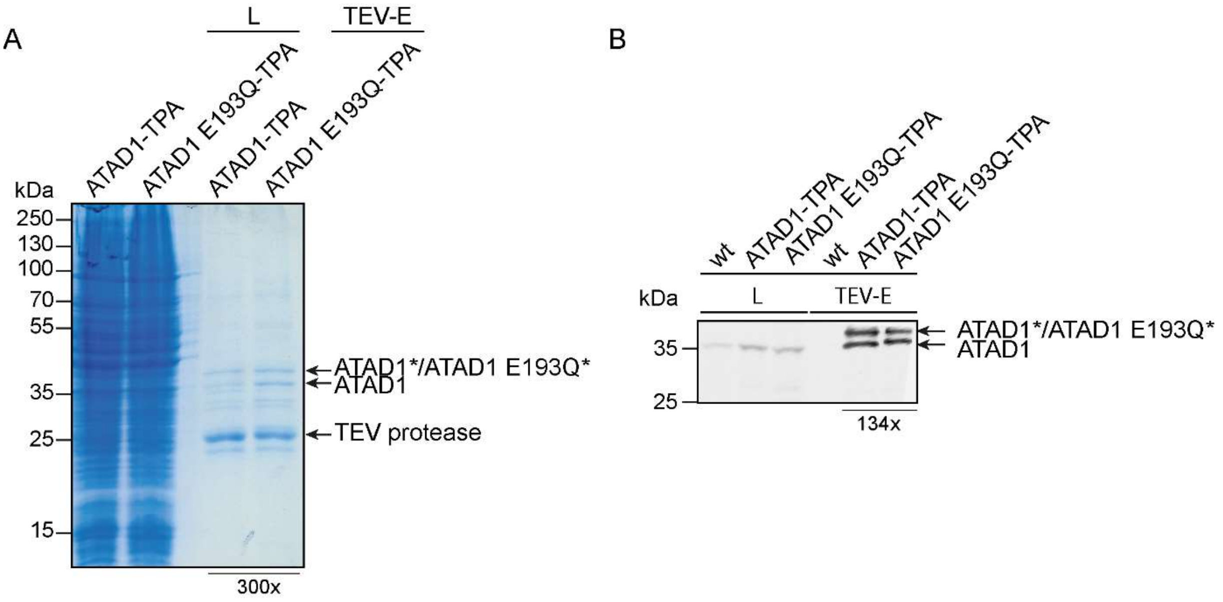


**Supplementary Fig. 2: Isolation of membrane-associated ATAD1-TPA and ATAD1 E193Q-TPA complexes out of human Flp-In293 cells. A:** Coomassie-stained gel shows the enrichment of several proteins by affinity purification of ATAD1-/ATAD1 E193Q-complexes. Besides TEV-cleaved ATAD1 which is denoted with an asterisk (*) also endogenous protein could be detected in the eluates. **B:** Immunoblot-staining with antibodies against ATAD1 shows the IgG affinity purification from Flp-In293 wild type cell line as negative control compared to the Flp-In ATAD1-TPA/ATAD1 E192Q-TPA cell lines. L: lysate, TEV-E: TEV-eluate.

**Supplementary Table 1: List of proteins found in ATAD1 and ATAD1 E193Q complexes.** Membrane-associated ATAD1-TPA and ATAD1 E193Q-TPA complexes were isolated from Flp-In293 cells and analysed by mass spectrometric analysis (n=5). As a negative control Flp-In293 cells without TPA fusion protein were used (n=2). Protein quantification was performed by the MaxLFQ algorithm. Proteins were found at least 4-times in the complex, were more than 5fold enriched in the ATAD1 eluate compared to the negative control eluates (LFQ ratio ATAD1(E193Q)/control) and had a p-value of <0.05. Proteins are listed in descending order of the LFQ value in ATAD1 complex. Some proteins were enriched in ATAD1 E193Q complexes compared to ATAD1 complexes (LFQ value ATAD1 E193Q/ATAD1 >1). The known localization according to UniProt as well as (at least partial) membrane-association are indicated. ER: endoplasmic reticulum; N: nucleus; M: mitochondria; P: peroxisomes; PM: plasma membrane.

| UniProt ID | Gene names | LFQ value (mean, n=5) | | LFQ ratio (n=2) | | Enrichment factor (LFQ value ATAD1 E193Q/ATAD1) | Localization (UniProt) | Membrane-associated (at least partially) |
|---|---|---|---|---|---|---|---|---|
| | | ATAD1 | ATAD1 E193Q | ATAD1/ control | ATAD1 E193Q/ control | | | |
| O43707 | ACTN4 | 2.19E+09 | 3.74E+08 | 23.2 | 4.0 | 0.17 | N, cytoskeleton (stress fiber), cytoplasm | |
| Q8NBU5 | ATAD1 | 2.11E+09 | 5.63E+09 | 211.0 | 562.2 | 2.67 | M, P, cytosol | x |
| P63261;P60709 | ACTG1;ACTB | 1.94E+09 | 1.01E+09 | 6.6 | 3.4 | 0.52 | cytoskeleton; cytoskeleton, N | |
| P21796 | VDAC1 | 1.17E+09 | 1.58E+09 | 8.6 | 11.6 | 1.35 | cell membrane, M | x |
| P35232 | PHB | 5.85E+08 | 2.06E+09 | 4.4 | 15.6 | 3.51 | M, N | x |
| P45880 | VDAC2 | 4.21E+08 | 9.25E+08 | 6.5 | 14.4 | 2.20 | M | x |
| P28288 | ABCD3 | 3.89E+08 | 1.25E+08 | 16.7 | 5.4 | 0.32 | P | x |
| P12236 | SLC25A6 | 3.72E+08 | 6.71E+08 | 3.0 | 5.5 | 1.81 | M | x |
| Q99623 | PHB2 | 3.63E+08 | 1.35E+09 | 4.4 | 16.5 | 3.71 | M, N, cytoplasm | x |
| P35580 | MYH10 | 3.19E+08 | 1.65E+08 | 5.4 | 2.8 | 0.52 | Lamellipodium | |
| Q8WVX9 | FAR1 | 2.87E+08 | 6.91E+07 | 9.7 | 2.3 | 0.24 | P | x |
| P05141 | SLC25A5 | 1.63E+08 | 3.16E+08 | 2.7 | 5.3 | 1.93 | M | x |
| Q9Y277 | VDAC3 | 1.58E+08 | 2.60E+08 | 7.4 | 12.2 | 1.65 | M | x |
| Q00325 | SLC25A3 | 1.36E+08 | 1.84E+08 | 3.7 | 5.0 | 1.35 | M | x |
| Q9H9B4 | SFXN1 | 1.32E+08 | 1.48E+08 | 4.7 | 5.3 | 1.13 | M | x |
| Q9Y6C9 | MTCH2 | 1.04E+08 | 1.34E+08 | 10.1 | 12.9 | 1.28 | M | x |
| O96011 | PEX11B | 9.62E+07 | 4.15E+07 | 73.4 | 31.7 | 0.43 | P | x |
| P12814 | ACTN1 | 9.38E+07 | 1.89E+07 | 7.3 | 1.5 | 0.20 | cell membrane, cytoskeleton | |
| Q9NR77 | PXMP2 | 8.95E+07 | 1.49E+07 | 56.5 | 9.4 | 0.17 | P | x |
| O00151 | PDLIM1 | 8.21E+07 | 5.82E+06 | 118.3 | 8.4 | 0.07 | cytoskeleton, cytoplasm | |
| P13073 | COX4I1 | 8.17E+07 | 1.74E+08 | 3.1 | 6.6 | 2.13 | M | x |
| O96008 | TOMM40 | 8.03E+07 | 1.01E+08 | 6.4 | 8.0 | 1.25 | M | x |
| Q02978 | SLC25A11 | 7.33E+07 | 1.29E+08 | 3.5 | 6.1 | 1.76 | M | x |
| Q9GZY8 | MFF | 6.82E+07 | 3.69E+07 | 49.1 | 26.6 | 0.54 | P, M | x |
| Q16643 | DBN1 | 4.52E+07 | 5.43E+07 | 6.1 | 7.3 | 1.20 | cytoplasm | |
| Q92614 | MYO18A | 4.22E+07 | 6.07E+07 | 89.2 | 128.2 | 1.44 | Golgi, cytoskeleton | |
| Q9UJS0 | SLC25A13 | 4.19E+07 | 8.99E+07 | 2.6 | 5.6 | 2.14 | M | x |
| Q9NS69 | TOMM22 | 3.82E+07 | 4.64E+07 | 5.7 | 6.9 | 1.21 | M | x |
| P00403 | MT-CO2 | 3.81E+07 | 1.14E+08 | 2.8 | 8.4 | 2.99 | M | x |
| P40855 | PEX19 | 3.80E+07 | 2.96E+07 | 19.8 | 15.5 | 0.78 | P, cytoplasm | x |

| UniProt ID | Gene names | LFQ value (mean, n=5) | | LFQ ratio (n=2) | | Enrichment factor (LFQ value ATAD1 E193Q/ATAD1) | Localization (UniProt) | Membrane-associated (at least partially) |
|---|---|---|---|---|---|---|---|---|
| | | ATAD1 | ATAD1 E193Q | ATAD1/ control | ATAD1 E193Q/ control | | | |
| Q8NBX0 | SCCPDH | 3.73E+07 | 1.84E+07 | 7.9 | 3.9 | 0.49 | extracellular, M, N, PM | |
| Q13608 | PEX6 | 3.57E+07 | 4.47E+07 | 6.3 | 7.9 | 1.25 | P, cytoplasm | x |
| P56589 | PEX3 | 3.50E+07 | 1.10E+07 | 39.9 | 12.5 | 0.31 | P | x |
| Q9UHQ9 | CYB5R1 | 3.41E+07 | 4.16E+07 | 8.8 | 10.8 | 1.22 | / | |
| Q8IXI2 | RHOT1/MIRO1 | 3.18E+07 | 2.62E+07 | 5.9 | 4.9 | 0.82 | M | x |
| O43933 | PEX1 | 3.07E+07 | 4.13E+07 | 5.5 | 7.4 | 1.34 | P, cytoplasm | x |
| P05413 | FABP3 | 2.96E+07 | 4.55E+05 | 125.1 | 1.9 | 0.02 | cytoplasm | |
| P53007 | SLC25A1 | 2.92E+07 | 4.14E+07 | 5.9 | 8.3 | 1.41 | M | x |
| O75083 | WDR1 | 2.92E+07 | 1.16E+06 | 35.2 | 1.4 | 0.04 | cytoskeleton | |
| O15228 | GNPAT | 2.90E+07 | 1.61E+07 | 5.5 | 3.0 | 0.56 | P | x |
| P10606 | COX5B | 2.81E+07 | 5.46E+07 | 2.9 | 5.6 | 1.94 | M | x |
| P33897 | ABCD1 | 2.80E+07 | 1.04E+07 | 6.5 | 2.4 | 0.37 | ER, Lysosome, M, P | x |
| Q9BWM7 | SFXN3 | 2.78E+07 | 4.20E+07 | 4.6 | 6.9 | 1.51 | M | x |
| Q96K12 | FAR2 | 2.77E+07 | 4.35E+06 | 9.8 | 1.5 | 0.16 | P | x |
| Q8IXI1 | RHOT2 | 2.65E+07 | 3.46E+07 | 6.4 | 8.4 | 1.31 | M | x |
| Q9NZ45 | CISD1 | 2.42E+07 | 1.07E+07 | 12.3 | 5.5 | 0.44 | M | x |
| Q7Z406 | MYH14 | 2.41E+07 | 7.37E+06 | 6.7 | 2.0 | 0.31 | cytoskeleton, cytosol, extracellular | |
| Q6P4A7 | SFXN4 | 2.38E+07 | 3.14E+07 | 10.3 | 13.7 | 1.32 | M | x |
| Q05682 | CALD1 | 2.33E+07 | 2.50E+07 | 40.2 | 43.1 | 1.07 | cytoskeleton, stress fiber | |
| Q9NX40 | OCIAD1 | 2.19E+07 | 5.68E+07 | 5.5 | 14.3 | 2.60 | endosome | |
| O43808 | SLC25A17 | 2.17E+07 | 4.12E+06 | 37.5 | 7.1 | 0.19 | P | x |
| O43676 | NDUFB3 | 2.14E+07 | 2.61E+07 | 6.1 | 7.4 | 1.22 | M | x |
| Q15388 | TOMM20 | 2.12E+07 | 1.85E+07 | 5.3 | 4.7 | 0.87 | M | x |
| P33121 | ACSL1 | 2.05E+07 | 1.18E+07 | 5.2 | 3.0 | 0.57 | M, P, ER | x |
| O14880 | MGST3 | 1.99E+07 | 1.62E+07 | 15.9 | 13.0 | 0.82 | ER | x |
| O95140 | MFN2 | 1.95E+07 | 1.91E+07 | 5.9 | 5.7 | 0.98 | M | x |
| Q96A26 | FAM162A | 1.80E+07 | 1.30E+07 | 6.0 | 4.4 | 0.72 | M | x |
| P57105 | SYNJ2BP | 1.75E+07 | 9.90E+06 | 8.8 | 5.0 | 0.56 | M | x |
| Q3SXM5 | HSDL1 | 1.75E+07 | 3.95E+06 | 9.8 | 2.2 | 0.23 | M | |
| P06858 | LPL | 1.68E+07 | 7.76E+05 | 39.9 | 1.8 | 0.05 | extracellular, PM | x |
| P50542 | PEX5 | 1.65E+07 | 4.89E+06 | 12.1 | 3.6 | 0.30 | P, cytoplasm | x |

| UniProt ID | Gene names | LFQ value (mean, n=5) | | LFQ ratio (n=2) | | Enrichment factor (LFQ value ATAD1 E193Q/ATAD1) | Localization (UniProt) | Membrane-associated (at least partially) |
|---|---|---|---|---|---|---|---|---|
| | | ATAD1 | ATAD1 E193Q | ATAD1/ control | ATAD1 E193Q/ control | | | |
| Q6NUK1 | SLC25A24 | 1.63E+07 | 4.22E+07 | 4.5 | 11.6 | 2.59 | M | x |
| P21397 | MAOA | 1.52E+07 | 1.32E+07 | 13.6 | 11.8 | 0.87 | M | x |
| Q9NZJ7 | MTCH1 | 1.48E+07 | 2.34E+07 | 8.4 | 13.3 | 1.58 | M | x |
| Q9NX47 | MARCH5 | 1.44E+07 | 1.83E+07 | 14.6 | 18.5 | 1.27 | ER, M | x |
| Q96KQ4 | PPP1R13B | 1.44E+07 | 2.04E+07 | 4.2 | 5.9 | 1.41 | cytoplasm, N | |
| Q969Z3 | MARC2 | 1.40E+07 | 2.28E+07 | 22.4 | 36.4 | 1.63 | M, P | x |
| P48739 | PITPNB | 1.38E+07 | 4.83E+06 | 5.3 | 1.9 | 0.35 | Golgi, cytoplasm | |
| P17152 | TMEM11 | 1.37E+07 | 1.16E+07 | 6.4 | 5.4 | 0.84 | M | x |
| Q7Z434 | MAVS | 1.34E+07 | 1.95E+07 | 4.1 | 6.0 | 1.46 | M, P | x |
| Q9Y5Y5 | PEX16 | 1.23E+07 | 6.63E+06 | 19.0 | 10.3 | 0.54 | P | x |
| P28328 | PEX2 | 1.20E+07 | 3.40E+06 | 14.0 | 4.0 | 0.28 | P | x |
| O60683 | PEX10 | 1.17E+07 | 4.18E+06 | 18.6 | 6.7 | 0.36 | P | x |
| P14406 | COX7A2 | 1.16E+07 | 2.54E+07 | 3.3 | 7.2 | 2.20 | M | x |
| Q53H12 | AGK | 1.13E+07 | 3.57E+07 | 1.9 | 5.9 | 3.15 | M | x |
| P12235 | SLC25A4 | 1.10E+07 | 2.14E+07 | 3.3 | 6.5 | 1.95 | M | x |
| Q9HD20 | ATP13A1 | 1.06E+07 | 4.80E+07 | 4.4 | 19.8 | 4.54 | ER | x |
| O75746 | SLC25A12 | 1.01E+07 | 2.20E+07 | 4.4 | 9.5 | 2.18 | M | x |
| Q9UH62 | ARMCX3 | 1.00E+07 | 1.40E+07 | 5.5 | 7.7 | 1.40 | M, N, cytoplasm | x |
| Q13232 | NME3 | 9.71E+06 | 1.09E+07 | 6.7 | 7.5 | 1.12 | cytosol | |
| O43491 | EPB41L2 | 8.97E+06 | 5.78E+06 | 10.1 | 6.5 | 0.64 | PM, cytoskeleton | |
| Q7KZN9 | COX15 | 8.78E+06 | 1.38E+07 | 6.1 | 9.7 | 1.57 | M | x |
| Q96ER9 | CCDC51 | 8.26E+06 | 2.83E+07 | 2.2 | 7.4 | 3.42 | M | x |
| P04899 | GNAI2 | 8.16E+06 | 3.06E+06 | 5.2 | 2.0 | 0.38 | PM, cytoskeleton, cytoplasm | |
| P67936 | TPM4 | 8.14E+06 | 4.21E+06 | 8.9 | 4.6 | 0.52 | cytoskeleton | |
| Q8N2F6 | ARMC10 | 7.92E+06 | 9.07E+06 | 18.2 | 20.9 | 1.15 | ER | x |
| O00623 | PEX12 | 7.91E+06 | 3.26E+06 | | | 0.41 | P | x |
| Q15070 | OXA1L | 7.84E+06 | 1.74E+07 | 3.4 | 7.6 | 2.22 | M | x |
| Q8IWA4 | MFN1 | 7.55E+06 | 4.79E+06 | 5.6 | 3.5 | 0.63 | M, cytoplasm | x |
| Q9P0J0 | NDUFA13 | 7.15E+06 | 7.60E+06 | | | 1.06 | M | x |
| P30519 | HMOX2 | 6.70E+06 | 6.61E+07 | 2.2 | 21.3 | 9.87 | ER | |
| Q5VT66 | MTARC1 | 6.38E+06 | 1.09E+07 | 3.7 | 6.4 | 1.70 | M | x |

| UniProt ID | Gene names | LFQ value (mean, n=5) | | LFQ ratio (n=2) | | Enrichment factor (LFQ value ATAD1 E193Q/ATAD1) | Localization (UniProt) | Membrane-associated (at least partially) |
|---|---|---|---|---|---|---|---|---|
| | | ATAD1 | ATAD1 E193Q | ATAD1/ control | ATAD1 E193Q/ control | | | |
| Q9HC21 | SLC25A19 | 6.34E+06 | 9.98E+06 | 8.4 | 13.3 | 1.57 | M | x |
| Q9Y6I8 | PXMP4 | 6.05E+06 | 5.48E+06 | | | 0.91 | P | x |
| Q9BQ95 | ECSIT | 5.76E+06 | 1.26E+07 | 3.0 | 6.5 | 2.18 | M, N, cytoplasm | |
| Q96HA9 | PEX11G | 5.72E+06 | 4.02E+06 | 23.5 | 16.5 | 0.70 | P | x |
| P51970 | NDUFA8 | 5.42E+06 | 8.64E+06 | | | 1.59 | M | x |
| Q16611 | BAK1 | 5.23E+06 | 4.38E+06 | 8.9 | 7.5 | 0.84 | M | x |
| Q86UT6 | NLRX1 | 4.68E+06 | 1.60E+07 | 4.6 | 15.8 | 3.41 | M | |
| Q8N2A8 | PLD6 | 4.55E+06 | 9.83E+06 | | | 2.16 | M | x |
| Q70CQ3 | USP30 | 4.37E+06 | 3.47E+06 | | | 0.79 | M | |
| Q9NX00 | TMEM160 | 4.33E+06 | 5.17E+06 | | | 1.19 | membrane | x |
| P61803 | DAD1 | 4.09E+06 | 5.97E+06 | | | 1.46 | ER | x |
| Q8NAN2 | FAM73A | 3.96E+06 | 5.65E+06 | 5.2 | 7.5 | 1.43 | M | x |
| Q86UB9 | TMEM135 | 3.90E+06 | 1.93E+06 | | | 0.49 | M, P | x |
| Q9Y619 | SLC25A15 | 3.85E+06 | 6.56E+06 | | | 1.70 | M | x |
| Q9H3K2 | GHITM | 3.64E+06 | 6.19E+06 | | | 1.70 | M | x |
| O43772 | SLC25A20 | 3.54E+06 | 6.26E+06 | | | 1.77 | M | x |
| Q96AG3 | SLC25A46 | 3.50E+06 | 4.79E+06 | | | 1.37 | M | x |
| P53701 | HCCS | 3.37E+06 | 3.53E+06 | 7.8 | 8.1 | 1.05 | M | |
| Q9Y512 | SAMM50 | 3.06E+06 | 1.70E+07 | 1.0 | 5.8 | 5.58 | M, cytoplasm | x |
| Q9H4I3 | TRABD | 2.98E+06 | 8.55E+06 | 3.1 | 8.9 | 2.87 | / | |
| Q9H936;Q9H1K4 | SLC25A22;SLC25A18 | 2.96E+06 | 6.31E+06 | 4.6 | 9.8 | 2.14 | M | x |
| P32189;Q14409 | GK;GK3P | 2.87E+06 | 8.26E+06 | 2.2 | 6.3 | 2.88 | M | x |
| Q9P0S9 | TMEM14C | 2.68E+06 | 2.10E+06 | 5.8 | 4.5 | 0.78 | M | x |
| O43920 | NDUFS5 | 2.65E+06 | 6.17E+06 | | | 2.33 | M | x |
| Q9BVV7 | TIMM21 | 2.44E+06 | 7.04E+06 | 2.8 | 8.1 | 2.89 | M | x |
| Q7Z412 | PEX26 | 2.36E+06 | 2.58E+06 | | | 1.09 | P | x |
| P31483 | TIA1 | 2.33E+06 | 3.55E+06 | | | 1.52 | N | |
| Q49A26 | GLYR1 | 2.32E+06 | 2.40E+06 | 5.4 | 5.5 | 1.03 | N | |
| Q08554 | DSC1 | 2.26E+06 | 3.07E+06 | | | 1.36 | PM | x |
| Q9BUB7 | TMEM70 | 2.23E+06 | 2.49E+06 | 7.0 | 7.8 | 1.12 | M | x |
| P61020 | RAB5B | 2.20E+06 | 2.48E+06 | | | 1.13 | Endosome, PM | x |

| UniProt ID | Gene names | LFQ value (mean, n=5) | | LFQ ratio (n=2) | | Enrichment factor (LFQ value ATAD1 E193Q/ATAD1) | Localization (UniProt) | Membrane-associated (at least partially) |
|---|---|---|---|---|---|---|---|---|
| | | ATAD1 | ATAD1 E193Q | ATAD1/ control | ATAD1 E193Q/ control | | | |
| P37840 | SNCA | 2.19E+06 | 0.00E+00 | 5.5 | | / | secreted, N, cytoplasm | |
| Q7Z7K6 | CENPV | 2.13E+06 | 8.66E+05 | 5.3 | 2.2 | 0.41 | cytoskeleton, N | |
| Q9BRR6 | ADPGK | 2.10E+06 | 2.06E+06 | | | 0.98 | secreted | |
| Q9BRX8 | FAM213A | 2.01E+06 | 1.10E+06 | 5.2 | 2.9 | 0.55 | secreted, cytoplasm | |
| O60830 | TIMM17B | 1.97E+06 | 5.54E+06 | | | 2.82 | M | x |
| Q969V5 | MUL1 | 1.93E+06 | 1.82E+06 | | | 0.94 | M, P | x |
| Q6IQ22 | RAB12 | 1.92E+06 | 2.37E+06 | | | 1.23 | Golgi, lysosome, endosome | x |
| Q5UIP0 | RIF1 | 1.89E+06 | 1.72E+06 | | | 0.91 | cytoskeleton, N | |
| Q14980 | NUMA1 | 1.83E+06 | 1.62E+06 | | | 0.88 | cytoskeleton, PM, N | |
| Q96BW9 | TAMM41 | 1.64E+06 | 3.10E+06 | 3.0 | 5.8 | 1.89 | M | |
| Q99665 | IL12RB2 | 1.58E+06 | 2.95E+06 | | | 1.87 | / | |
| Q96D53 | ADCK4 | 1.49E+06 | 7.75E+05 | 5.2 | 2.7 | 0.52 | cytosol, PM, M | x |
| Q9BQE5 | APOL2 | 1.39E+06 | 1.31E+06 | 5.8 | 5.5 | 0.95 | / | |
| Q15334 | LLGL1 | 1.30E+06 | 8.55E+05 | | | 0.66 | Golgi, endosome, cytoskeleton | |
| Q9UHB6 | LIMA1 | 1.17E+06 | 1.45E+07 | | | 12.43 | cytoskeleton, PM | |
| Q9BUR5 | APOO | 1.01E+06 | 5.30E+06 | | | 5.23 | Golgi, ER, M | x |
| P50851 | LRBA | 9.77E+05 | 6.85E+05 | | | 0.70 | Golgi, lysosome, ER, PM | x |
| A6NKD9 | CCDC85C | 8.58E+05 | 1.19E+06 | 3.6 | 5.0 | 1.39 | / | |
| O75155 | CAND2 | 8.39E+05 | 6.54E+05 | | | 0.78 | N | |

**Supplementary Table 2: Mitochondrial proteins found in ATAD1 and ATAD1 E193Q complexes.** To show enrichment in wildtype or mutated complex the enrichment factor was calculated by dividing the LFQ-intensity through the negative control: LFQ ATAD1(E193Q)/LFQ control. Proteins marked with an asterisk were not observed in the control.

| protein | enrichment factor | | protein | enrichment factor | |
|---|---|---|---|---|---|
| | ATAD1 | ATAD1 E193Q | | ATAD1 | ATAD1 E193Q |
| MARCH5 | 14.6 | 18.5 | MTARC1 | 3.7 | 6.4 |
| PHB2 | 4.4 | 16.5 | GK | 2.2 | 6.3 |
| NLRX1 | 4.6 | 15.8 | SLC25A11 | 3.5 | 6.1 |
| PHB | 4.4 | 15.6 | AGK | 1.9 | 5.9 |
| VDAC2 | 6.5 | 14.4 | SAMM50 | 1.0 | 5.8 |
| SFXN4 | 10.3 | 13.7 | TAMM41 | 3.0 | 5.8 |
| SLC25A19 | 8.4 | 13.3 | MFN2 | 5.9 | 5.7 |
| MTCH1 | 8.4 | 13.3 | SLC25A13 | 2.6 | 5.6 |
| MTCH2 | 10.1 | 12.9 | COX5B | 2.9 | 5.6 |
| VDAC3 | 7.4 | 12.2 | CISD1 | 12.3 | 5.5 |
| MAOA | 13.6 | 11.8 | SLC25A6 | 3.0 | 5.5 |
| SLC25A24 | 4.5 | 11.6 | TMEM11 | 6.4 | 5.4 |
| VDAC1 | 8.6 | 11.6 | SFXN1 | 4.7 | 5.3 |
| SLC25A22 | 4.6 | 9.8 | SLC25A5 | 2.7 | 5.3 |
| COX15 | 6.1 | 9.7 | SLC25A3 | 3.7 | 5.0 |
| SLC25A12 | 4.4 | 9.5 | SYNJ2BP | 8.8 | 5.0 |
| MT-CO2 | 2.8 | 8.4 | RHOT1 | 5.9 | 4.9 |
| RHOT2 | 6.4 | 8.4 | TOMM20 | 5.3 | 4.7 |
| SLC25A1 | 5.9 | 8.3 | TMEM14C | 5.8 | 4.5 |
| HCCS | 7.8 | 8.1 | FAM162A | 6.0 | 4.4 |
| TIMM21 | 2.8 | 8.1 | SCCPDH | 7.9 | 3.9 |
| TOMM40 | 6.4 | 8.0 | MFN1 | 5.6 | 3.5 |
| TMEM70 | 7.0 | 7.8 | ADCK4 | 5.2 | 2.7 |
| ARMCX3 | 5.5 | 7.7 | HSDL1 | 9.8 | 2.2 |
| OXA1L | 3.4 | 7.6 | NDUFA13 | * | * |
| FAM73A | 5.2 | 7.5 | NDUFA8 | * | * |
| BAK1 | 8.9 | 7.5 | PLD6 | * | * |
| NDUFB3 | 6.1 | 7.4 | USP30 | * | * |
| CCDC51 | 2.2 | 7.4 | SLC25A15 | * | * |
| COX7A2 | 3.3 | 7.2 | GHITM | * | * |
| TOMM22 | 5.7 | 6.9 | SLC25A20 | * | * |
| SFXN3 | 4.6 | 6.9 | SLC25A46 | * | * |
| COX4I1 | 3.1 | 6.6 | NDUFS5 | * | * |
| SLC25A4 | 3.3 | 6.5 | TIMM17B | * | * |
| ECSIT | 3.0 | 6.5 | APOO | * | * |
